# Endothelial TET2 Regulates Cardiac Remodeling by Modifying Endothelial-to-Mesenchymal Transition

**DOI:** 10.1101/2022.06.15.496224

**Authors:** Wenxin Kou, Yefei Shi, Bo Li, Yanxi Zeng, Ming Zhai, Shuangjie You, Qing Yu, Shiyu Gong, Jianhui Zhuang, Yifan Zhao, Weixia Jian, Yawei Xu, Wenhui Peng

## Abstract

DNA methylation modification has been proved to play an important role in heart diseases. In this study, the role of Ten-Eleven Translocation-2 (TET2), which is a key demethylation enzyme, is investigated in cardiac remodeling. TET2 is abundant in endothelial cells but decreased in hypertrophic hearts. TET2 knockdown in endothelial cells triggers endothelial-to-mesenchymal transition (EndMT), while overexpression of TET2 inhibits the EndMT. In vivo, Cdh5-CreERT2/TET2^flox/flox^; Rosa26-mTmG^+/-^ mice are developed and undergo transverse aortic constriction (TAC) subsequently to induce pathological cardiac hypertrophy model. Hearts of Cdh5-CreERT2/TET2^flox/flox^ mice show more severe hypertrophy and fibrosis than controls in the TAC model. Furthermore, EGLN3 is identified to participate in EndMT as the downstream target of TET2 by using RNA sequencing and whole-genome bisulfite sequencing (WGBS). Finally, vitamin C, which can mimic TET2 restoration, is found to partially improve cardiac function and inhibit myocardial fibrosis. These insights into how TET2 alleviates cardiac fibrosis may open new avenues for treating cardiac remodeling in the future.

## 1. Introduction

Heart failure development is characterized by the process of adverse cardiac remodeling. Cardiac remodeling is a dynamic change of cardiac morphology and function when the heart undergoes different physiological and pathological stimulation^[1]^. One of the typical characteristics of pathological remodeling is cardiac fibrosis^[2]^. Usually, cardiac fibrosis will lead to heart stiffening and cause heart failure with preserved ejection fraction (HFpEF) in the early stage and reduced ejection fraction (HFrEF) in the advanced stage^[3]^. Cardiac fibrotic is a process achieved by a collective contribution from fibroblast, myofibroblast, and endothelial-to-mesenchymal transition (EndMT)^[4]^. EndMT refers to the pathological feature of endothelial cells losing initial endothelial characteristics and expressing typical markers of fibroblast^[5]^. It is acknowledged that EndMT regulates embryonic development of heart, such as atrioventricular valvuloseptal morphogenesis^[6]^. Recently, EndMT is reported to play an essential role in cardiac fibrosis and remodeling^[7, 8]^. Yet, its mechanism in cardiac remodeling is not clear so far.

Methyl group from S-adenyl methionine transferred to the fifth carbon of a cytosine residue to form 5-methylcytosine (5-mC) is called DNA methylation^[9]^. Studies have shown that DNA methylation is essential in the pathogenesis of fibrosis, including the progress of EndMT^[10]^. Ten-eleven translocation methylcytosine dioxygenase (TET) enzymes can catalyze 5-mC to 5-hydroxymethylcytosine (5-hmC), 5-formylcytosine (5-fC) or 5-carboxylcytosine (5-caC)^[11]^, resuming gene expression. Mounting evidence has confirmed the important roles of TET2 in cardiovascular diseases. Our previous study found that disequilibrium of DNA methylation caused by TET2 gene methylation in smooth muscle cells was important for vascular remodeling^[12]^. Clonal hematopoiesis, such as mutation of TET2 or Dnmt3, also influenced the vascular remodeling^[13]^. The abnormal function of hematopoietic stem/progenitor in clonal hematopoiesis would accelerate cardiac dysfunction and fibrosis^[14]^.

In our study, we explored the role of TET2 in cardiac fibrosis and EndMT, and found the loss of TET2 could promote EndMT and cardiac fibrosis. Cdh5-specific TET2 knockout lineage tracking mice were constructed to confirm its impact in vivo. Genome-wide RNA sequencing and whole-genome bisulfite sequencing (WGBS) were performed on human umbilical vein endothelial cells (HUVECs) treated by TGF-β1 and IL-1β with or without TET2 knocked down to explore the downstream genes. Finally, vitamin C, which could induce Tet-dependent DNA demethylation, reduced the EndMT and could be an adjuvant therapy for cardiac fibrosis.

## 2. Methods

### Human sample collection and ethical approval

Our study was approved by the Ethical Committee of Shanghai Tenth People’s Hospital, and our clinical experiments were complied with the relevant ethical regulations (Shanghai, China; permit number: 2020-KN50-01). Human heart tissues were obtained from autopsies. The written informed consents were collected from patients or their relatives.

### Animals

Tet2-flox mice (Tet2^tm1.1laai^/J) were purchased from The Jackson Laboratory, Cdh5-CreERT2, Rosa26-mTmG and *db/db* mice were purchased from Biocytogen (China). Mice were crossbreed to generate Cdh5-Cre/Tet2^-/-^; Rosa26-mTmG and Cdh5-Cre/Tet2^flox/flox^; Rosa26-mTmG. Tamoxifen (75 mg/kg body weight, Cat#T5648, Sigma, USA) was administrated by intraperitoneal injection every day for a total of five times to induce conditional activation of Cre recombinase. All the animal experimental operations were performed following the guidelines of the National Institutes of Health for the care and use of laboratory animals (NIH Publication, 8th Edition, 2011) and all the experimental procedures involving animals were approved by the Animal Care and Use Committees of Shanghai Tenth People’s Hospital (Shanghai, China; permit number: SHDSYY-2020-2149).

### Transverse aortic constriction

Male mice of 6-8 weeks of age were used for pathological cardiac remodeling studies. Mice were anesthetized with pentobarbital sodium (50 mg/kg, Intraperitoneal injection). Then the chest of mice was opened with the minimally invasive method. The transverse aorta between the right innominate and left common carotid arteries was ligated with a 27-gauge needle by a 6–0 silk suture to form an approximately 70% aortic constriction. The needle was then gently removed, and the thoracic cavity was closed. The sham procedure was identical except for the partial aorta ligation.

### Myocardial infarction model

Myocardial infarction was performed on mice. Briefly, thoracotomy was conducted in the fourth left intercostal space after anaesthetization. The thorax and pericardium were opened, and the left anterior descending coronary artery was ligated with a needle with 7-0 silk suture. Mice with sham procedure served as controls.

### Cell culture and induction of EndMT

Human umbilical vein endothelial cells (HUVECs, Cat#8000, ScienCell, USA) were cultured at 37 °C with 5% CO_2_ in endothelial cells medium (ECM, Cat#1001, ScienCell, USA) with 5% fetal bovine serum (FBS, Cat#0025, ScienCell, USA), 1% endothelial cell growth supplement (ECGS, Cat#1052, ScienCell, USA) and 1% penicillin/streptomycin solution (P/S, Cat#0503, ScienCell, USA). To induce EndMT, HUVECs were cultured in a complete growth medium until 50% confluence. The medium was changed for complete growth medium containing TGF-β (10 ng/ml, Cat#100-21, PeproTech, USA) and IL-1β (1ng/ml, Cat#200-01B, PeproTech, USA) for five days.

### Primary cells isolation

Neonatal mouse cardiomyocytes, fibroblasts, and lung endothelial cells were isolated as previously described^[15]^. In brief, ventricles of hearts were removed from 3-day-old C57BL/6J mice and finely minced followed by sequential digestion with 1 mg/ml collagenase II (Cat#1148090, Sigma). The collected primary cells were screened through a 70 μm cell strainer and seeded onto uncoated 6-well cell culture plates in DMEM medium containing 20% FBS and 1% P/S. After culturing for 60 min at 37 °C, the supernatant (containing the cardiomyocytes) was seeded onto FBS-coated 6-well cell culture plates, and the adherent cells (almost exclusively primary cardiac fibroblasts) were cultured in DMEM with 15% FBS and 1% P/S. Mouse lung endothelial cells were isolated with CD31-positive immunomagnetic beads. Briefly, lungs were removed from 3-day-old C57BL/6J mice and mechanically minced followed by digestion with 2 mg/ml collagenase II at 37 °C for 40 min. After filtration using sterile 70 μm strainers, the cells were incubated with the anti-CD 31 antibody (Cat# 557355, BD biosciences) at 4 °C overnight. The next day, cells were incubated with dynabeads at 4 °C for 45 min and magnetic separation of the dynabeads from the supernatant.

### SiRNA Transfection and adenovirus transduction

TET2 siRNA and control siRNA (50 nM) were transfected into HUVECs using Lipofectamine 2000 (Cat#11668019, Invitrogen, USA) following the manufacturer’s instructions. The sense sequence (5’-3’) of TET2 is GCUAAAUACCUGUUCCUUUTT, and the antisense sequence (5’-3’) of TET2 is AAAGGAACAGGUAUUUAGCTT. The sense sequence (5’-3’) of EGLN3 is GCAACCAGAUAUGCUAUGATT, and the antisense sequence (5’-3’) of TET2 is UCAUAGCAUAUCUGGUUGCTT. The sense sequence (5’-3’) of negative control is UUCUCCGAACGUGUCACGUTT, and the antisense sequence (5’-3’) of negative control is ACGUGACACGUUCGGAGAATT.

TET2 adenovirus and control adenovirus particles were produced by Shanghai Genechem Co. Ltd. HUVECs were incubated with adenovirus at a multiplicity of infection (MOI) of 50 overnight in complete growth medium. We replaced the complete growth medium with fresh complete medium the following day.

### RNA isolation and quantitative polymerase chain reaction

Total RNA from cultured cells was isolated using TRIzol reagent (Cat#15596026, Invitrogen, USA) according to the manufacturer’s instruction. 1 µg of total RNA was used for reverse transcription using PrimeScript RT Reagent Kit (Cat#RR037A, TaKaRa, Japan). Quantitative polymerase chain reaction (qPCR) was performed using KAPA SYBR FAST qPCR Master Mix (KM4101, KAPA Biosystems, USA). The 2^-ΔΔCT^ method was used for quantitative analysis. The sequence of primers is presented in Supplementary Table 1.

### Western blot analysis

Cultured cells were lysed in Cell Lysis Buffer (Cat#9803s, Cell Signaling Technology, USA) with protease inhibitors (Cat#04693159001, Roche, Switzerland) for 20 min on ice. After centrifugation at the speed of 12,000 g for 10 min at 4 °C, the protein content of the samples was determined using the bicinchoninic acid kit (Cat#20201ES76, Yeasen, China). Equal amounts of protein were loaded onto SDS-polyacrylamide gels and transferred to PVDF membranes. After being blocked by 5% bovine serum albumin for 1 h, membranes were incubated with primary antibodies at 4 °C overnight. The antibodies’ information and titers are presented in Supplementary Table 2. The PVDF membranes were washed in PBST and then incubated with horseradish peroxidase-conjugated secondary antibodies. Bands were visualized using chemiluminescence (Cat#180-5001; Tanon, China), detected by Amersham Imager 600 system (GE Healthcare Life Sciences, USA), and quantitated by ImageJ software (v1.52a, National Institutes of Health, USA).

### EdU assay

After starvation for 6 h, HUVECs were incubated with 10 μM EdU (5-ethynyl-2′-deoxyuridine) for 12 h. Detection of EdU was performed using Click-iT(tm) EdU Cell Proliferation Kit (Cat#10339, Invitrogen, USA).

### Migration assay

When HUVECs reached 90% confluence in 6-well culture plates, the medium was replaced by ECM with 0.5% FBS. After starvation for 12 h, a 200 μl pipette was used to make a wound across the well. Pictures were captured at the time of 0 h, 12 h, 24h after scratching.

### DNA isolation and DNA methylation analysis

Global DNA from cultured cells was isolated using TIANamp Genomic DNA Kit (Cat#DP304, Qiangen, China) according to the manufacturer’s instruction. A Global DNA Methylation Assay Kit (Cat#55017, Active Motif, USA) was used for DNA methylation analysis. 1 µg of global DNA was isolated to quantify the DNA methylation according to the manufacturer’s instruction.

### Dot blot

Cellular DNA was isolated using TIANamp Genomic DNA Kit (Cat#DP304, Tiangen, China) according to the manufacturer’s instruction. DNA was blotted onto a nitrocellulose membrane and incubated with anti-5-hmC at 4°C overnight. After the incubation with horseradish peroxidase-conjugated secondary antibody, the signal was visualized with chemiluminescence (Cat#180-5001, Tanon, China). Methylene blue staining of the membranes was used to assess equal DNA loading.

### Immunohistochemistry

Paraffin-embedded tissue sections were treated with microwave-based antigen retrieval using 10 mM sodium citrate buffer and then incubated with 0.3% hydrogen peroxide for 10 min. After permeabilization with 0.2% Triton X-100 for 15 min, sections were incubated at 4 °C overnight with primary antibodies. After incubation with secondary antibodies and amplification with streptavidin-biotin, sections were visulized using a DAB peroxidase substrate kit (Vector Laboratories, USA) and counterstained with hematoxylin.

### Immunofluorescent staining

Frozen sections were incubated with 4% paraformaldehyde for 15 min, followed by permeabilization with 0.2% Triton X-100 in PBS for 15 min. Then the sections were incubated with primary antibodies listed in Supplementary Table 2 at 4 °C overnight. After washing with PBS, secondary antibodies were incubated for 1 h at 37 °C in the dark, and the nuclei were stained with DAPI (1:5000, Vector Laboratories). Fluorescence images were obtained by an Olympus IX83 fluorescence microscope (Olympus Corporation, Japan).

### High-throughput RNA sequencing

HUVECs were transfected with siRNA (siTET2 or negative control, 50 nM) for 48 h. After starvation for 12 h, TGF-β (10 ng/mL) and IL-1β (1ng/mL) were used to stimulate HUVECs for another five days. Total RNA was isolated as previously described using Trizol (Cat# 15596026, Thermo Fisher) and extracted by chloroform and isopropanol^[16]^. RNA sequencing was conducted with the help of Beijing Berry Genomics Co.,Ltd. We prepared RNA libraries for sequencing using standard Illumina protocols, and subsequently performed RNA sequencing using the Illumina novaseq 6000 platform. Finally, differentially expressed genes were screened using DESeq (thresholds: linear fold change >1 or < -1, P < 0.05). Raw and processed data were deposited in GEO (GSE195696).

### Whole-genome bisulfite sequencing

Global DNA from cultured cells was isolated using TIANamp Genomic DNA Kit, and then fragmented by sonication to a mean size of approximate 250 bp. Next, genomic DNA library was constructed and ligated DNA was bisulfite-converted using the EZ DNA Methylation-Gold kit. Sequencing was performed using the Illumina Hiseq 4000 platform (Illumina, San Diego, CA, USA). After filtering, WGBS analyses were conducted by OE Biotech Co.,Ltd. (Shanghai, China). The Fisher test was applied to detect significant DMRs, and the screened criteria as methylation differences > 10%, P value < 0.05. After DMRs were identified, differentially methylated genes (DMGs) located in DMRs were characterized. Raw and processed data were deposited in GEO (GSE195696).

### Functional enrichment analysis of differentially expressed genes

To extract functional enrichment analysis, including biological processes, ellular components and molecular functions, and KEGG pathway analysis of the overlapping DEGs, DAVID (https://david.abcc.ncifcrf.gov/) was used, which is an online bioinformatics tool composed of a comprehensive biological knowledge base and analytic tool. The bubble chart was plotted by http://www.bioinformatics.com.cn, an online platform for data analysis and visualization.

### Enzyme-linked immunosorbent assay

Enzyme-linked immunosorbent assay (ELISA) was carried out using commercially available kits in accordance with the manufacturer’s instructions. TGF-β ELISA kit was obtained from Sino Biological Inc (Cat#KIT10804). Briefly, analytes were added into 96-well plates pre-coated with the antibody and incubated at 37 °C. After 2 h, the liquid was discarded and washed completely. Next, substrate solution was added to each well and incubated at 37 °C for 20 minutes. Finally, a stop solution was infused into each well, and absorbance was measured at 450 nm.

### Flow cytometry

Mice were intracardially perfused with 10 mL of PBS after anesthetization. The hearts were dissected, minced with fine scissors, and enzymatically digested in Hanks’ Balanced Salt Solution (HBSS) with 2 mg/mL collagenase II, 100U/mL DNase I and hyaluronidase at 37 °C for 1 h with gentle agitation. After digestion, the samples were filtered through a 70 µm strainer and incubated in the red blood cell lysis for 15 min. The obtained cells were washed with HBSS and stained with antibodies, including APC anti-mouse vimentin. Cells were sorted using FACS Aria III (BD Biosciences), and analyzed with FlowJo V10.7.1.

### Statistical analysis

Data were expressed as mean ± SEM. Comparisons between two groups were calculated using student’s *t*-test. The difference across three or more groups were tested by one-way ANOVA followed by a post hoc analysis with Bonferroni test. *P*-value < 0.05 was considered to indicate statistical significance. Statistical analyses were performed using Graphpad Prism 8.0.

## 3. Results

### 3.1 TET2 expression is decreased in cardiac remodeling

Various studies has demonstrated that DNA methylation could regulate heart failure, while its mechanism has not been fully understood^[17]^. Therefore, it is important to study the role of DNA methylation in the change of cardiac structure and function and how to alleviate cardiac remodeling in the early stage by regulating DNA methylation levels. As 5-azacitidine (5-aza) can drive DNA hypomethylation^[18]^, we injected 5-aza intraperitoneally in C57BL/6J mice undergoing TAC. Echocardiography was used to evaluate heart function. We observed that the cardiac function was better in mice treated with 5-aza compared with controls (Suppl. Figure 1a). Specifically, ejection fraction (EF) in the TAC group significantly increased from 39% to 50% after 5-aza injection (p = 0.04), with decreased left ventricular mass (LV mass), left ventricular end-systolic inner diameter (LVID; s) and left ventricular systolic posterior wall thickness (LVPW; s) (Suppl. Figure 1b). Moreover, HE and Masson’s trichrome staining showed that TAC challenge led to disarranged cardiomyocytes and severe fibrosis, which was alleviated to 41% by 5-aza treatment (*p* = 0.0002) (Suppl. Figure 1c-1g). 5-aza also suppressed the expression of ANP and TGF-β in the TAC hearts (Suppl. Figure 1h). Collectively, these results demonstrated that DNA demethylation could inhibit cardiac hypertrophy. It has been previously reported that three mammalian TET enzymes (TET1, TET2, and TET3) all could promote DNA demethylation^[19]^. Thus, we next explored whether there is a tissue-specific role of these TETs by investigating the expression of TET 1-3 in hearts. Although the expression of all three TETs decreased, the change in TET2 was the most significant (Suppl. Figure 2), and as such, we chose TET2 as our target for further study.

Western blot and immunohistochemistry demonstrated that TET2 expression was significantly decreased after TAC compared with sham group (Figure 1a-b, 1d). Similar results were found in human hypertrophic heart (Figure 1c), further supporting the potential role of TET2 in cardiac remodeling. The next question was whether such trend is a general phenomenon during cardiac remodeling or is pressure-overload-condition specific. Hence, we checked the TET2 expression and translation levels in other cardiac remodeling models, including *db/db* mice and mice with myocardial infarction. We found that TET2 expression in the heart of *db/db* mice (Suppl. Figure 3a-3c) and myocardial infarction (Suppl. Figure 3d-3f) decreased compared with controls. We further examined the levels of TET2 in primary cardiomyocytes, cardiac endothelial cells, and primary embryonic fibroblasts, the three major cell types of heart. The expression of TET2 in endothelial cells was much higher than that in the other two types of cell (Figure 1g, 1j). Consistently, immunofluorescence staining confirmed that most TET2-positive cells merged with cells expressed CD31 (Figure 1h). Additionally, HUVECs treated with angiotensin II had a significantly decreased TET2 expression (Figure 1i, 1k). These results indicated that TET2 was down-regulated in hypertrophic hearts and was mainly expressed in endothelial cells in heart.

**Figure 1.**
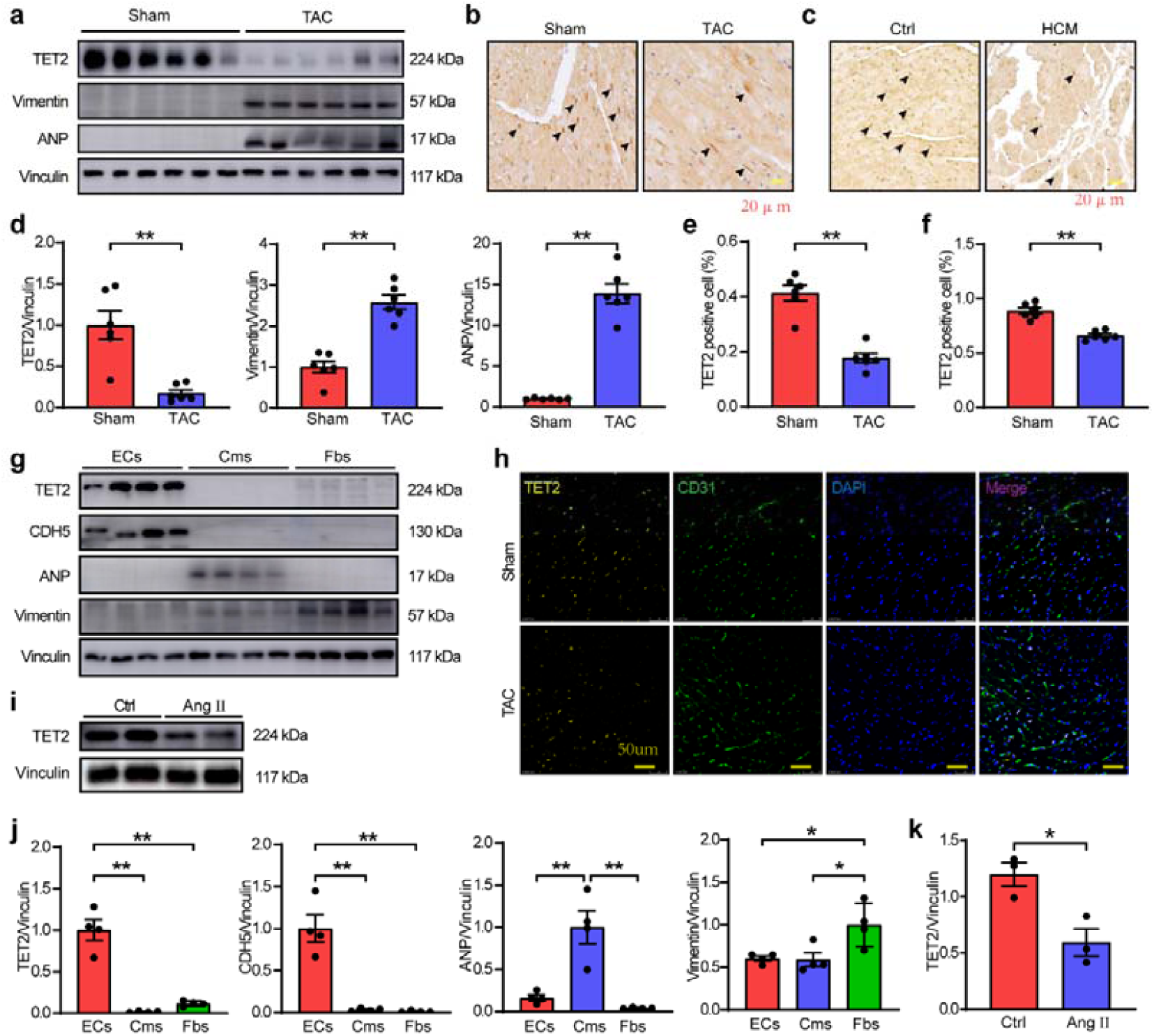
TET2 expression is decreased in cardiac remodeling. (a) The protein of TET2 decreased in TAC (n=6). (b) Immunohistochemistry of TET2 in sham and TAC mice (n=6). (c) Immunohistochemistry of TET2 in control and myocardial hypertrophy patients (n=6). (d) Statistical results of western blot in a. (e) Statistical results of b. (f) Statistical results of c. (g) The levels of TET2 in primary myocardial cells, cardiac endothelial cells, and primary embryonic fibroblasts (n=4). (h) Immunofluorescence staining of TET2 and CD31 in heart (n=5). (i) Western blot of HUVECs stimulated by Ang II (n=3). (j) Statistical results of g. (k) Statistical results of i. Data are expressed as mean ± SEM. *P<0.05, **P<0.01.

### 3.2 TET2 downregulation in HUVECs promotes endothelial-to-mesenchymal transition

The colocalization of TET2 expression with CD31 in hearts suggested that TET2 exerted its biological function mainly in endothelial cells. We then knocked down TET2 in HUVECs to explore the impact of TET2 on endothelial cells. HUVEC underwent morphology transformation from oval to fusiform after transfection (Figure 2a). In parallel with the morphological chance, the skewness of protein expression from endothelial marker VE-Cadherin dominant staged to mesenchymal marker vimentin expressing Figure 2b). The expression of endothelial markers decreased by 50%, while mesenchymal markers increased to 2.8 folds change after TET2 knocked down (p < 0.05), which suggested the deficiency of TET2 on HUVECs contributed to EndMT (Figure 2d). Since TGF-β can induce EndMT of ECs through activating receptor-regulated Smad (R-Smad)^[20]^, we further investigate the TGF-β/Smad pathway in TET2 knockdown in HUVECs and found the pathway activation upon TET2 deficiency. In addition, the concentrations of TGF-β increased about three times in the supernatant of siTET2 HUVECs compared to the control group (Figure 2c, 2f). ELISA and dot blot were conducted to test whether TET2 functioned via regulating DNA methylation. As expected, siTET2 HUVECs had higher levels of DNA methylation (Figure 2g, 2h), accompanied with aggravated HUVECs proliferation but unaffected migration (Suppl. Figure 4a-4d).

**Figure 2.**
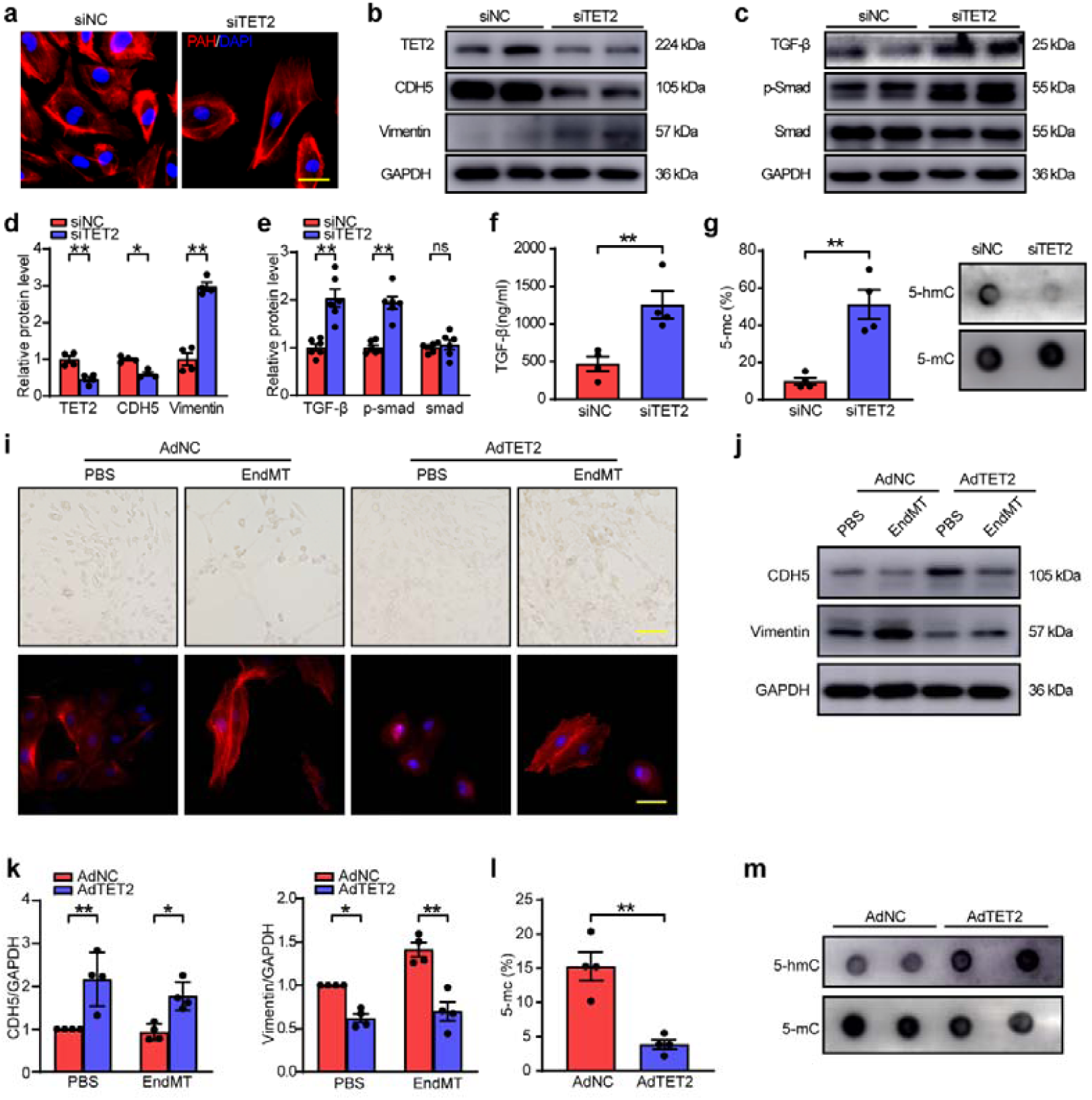
TET2 knocked down in HUVECs promotes endothelial-to-mesenchymal transition. (a) The morphology of HUVECs changed after TET2 knockdown. (b) Western blot of endothelial marker VE-Cadherin and mesenchymal marker vimentin in HUVECs after TET2 knockdown (n=4). (c) Western blot of TGF-β /Smad signaling in HUVECs after TET2 knockdown (n=6). (d) Statistical results of b. (e) Statistical results of c. (f) ELISA of TGF-β in the supernatant of siTET2 HUVECs (n=4). (g) ELISA of 5-mc in siTET2 HUVECs (n=4). (h) Dot blot showed higher levels of DNA methylation in siTET2 HUVECs (n=5). (i) The morphology of HUVECs undergoing EndMT with TET2 overexpressed. (j) Western blot of endothelial marker VE-Cadherin and mesenchymal marker vimentin in HUVECs after TET2 overexpression (n=4). (k) Statistical results of j. (l) ELISA of 5-mc in adTET2 HUVECs (n=4). (h) Dot blot showed higher levels of DNA methylation in adTET2 HUVECs (n=4). Data are expressed as mean ± SEM. *P<0.05, **P<0.01.

The role of TET2 was further analyzed in EndMT HUVECs. First, EndMT cell model was successfully established by validating cell morphology (Suppl. Figure 5a) and corresponding cell markers (Suppl. Figure 5b-5c, 5e-5f). DNA methylation levels were found to increase in EndMT HUVECs (Suppl. Figure 5g), consistent with our findings in siTET2 HUVECs. And the application of 5-aza could alleviate EndMT (Suppl. Figure 5h-5j). Consequently, by utilizing an adenoviral expression vector of TET2 construct (Suppl. Figure 6a) to overexpress TET2 in HUVEC, we found that the EndMT was significantly reduced (Figure 2i-2k) with less DNA methylation level (Figure 2l-2m) and proliferation (Suppl. Figure 6b-6e), in line with findings of the previous study^[11]^.

### 3.3 Endothelial cell-specific TET2 knockout aggravates cardiac function and fibrosis in pressure overload induced cardiac remodeling in vivo

Regeneration lineage tracing was performed by Cdh5-CreERT2 mice carrying the mTmG reporter^[21]^ (Suppl. Figure 7a) to explore the role of TET2 on endothelial cells in vivo and define EndMT contribution to fibrosis. Then we conditionally deleted TET2 in endothelial cells by crossing the TET2^flox/flox^ mice with Cdh5-CreERT2^+/−^ mice.

In Cdh5-CreERT2^+/-^; Rosa26-mTmG^+/-^ mice, fluorescence of cells expressing Cdh5 changed from tomato positive to GFP positive after tamoxifen injection (Suppl. Figure 7b). About 20% cardiac cells were GFP positive in an untreated Cdh5-CreERT2^+/-^; Rosa26-mTmG^+/-^ (WT) mouse (Suppl. Figure 7c). Moreover, analysis of the TET2 mRNA expression in Cdh5-CreERT2/TET2^flox/flox^; Rosa26-mTmG^+/-^ (TET2-CKO) hearts also confirmed that the TET2^flox/flox^ allele was efficiently recombined in the endothelial cells. GFP positive cells were sorted out by flow cytometry from TET2-CKO mice had only 30%-40% levels of TET2 compared to that of WT (Suppl. Figure 8).

TAC was performed in these mice. The heart function was worsened in the TET2-CKO mice compared with control mice based on multiple echocardiographic measurements (Figure 3a, 3b, 3e). Besides, cardiomyocyte size increased moderately in the TET2-CKO group. However, the degree of cardiac fibrosis in TET2-CKO was up to 2.5 times as much as that of WT mice (p < 0.01) (Figure 3c, 3d), suggesting that the deficiency of TET2 in endothelial cells exhibited a more significant influence on fibroblasts than cardiomyocytes in vivo.

**Figure 3.**
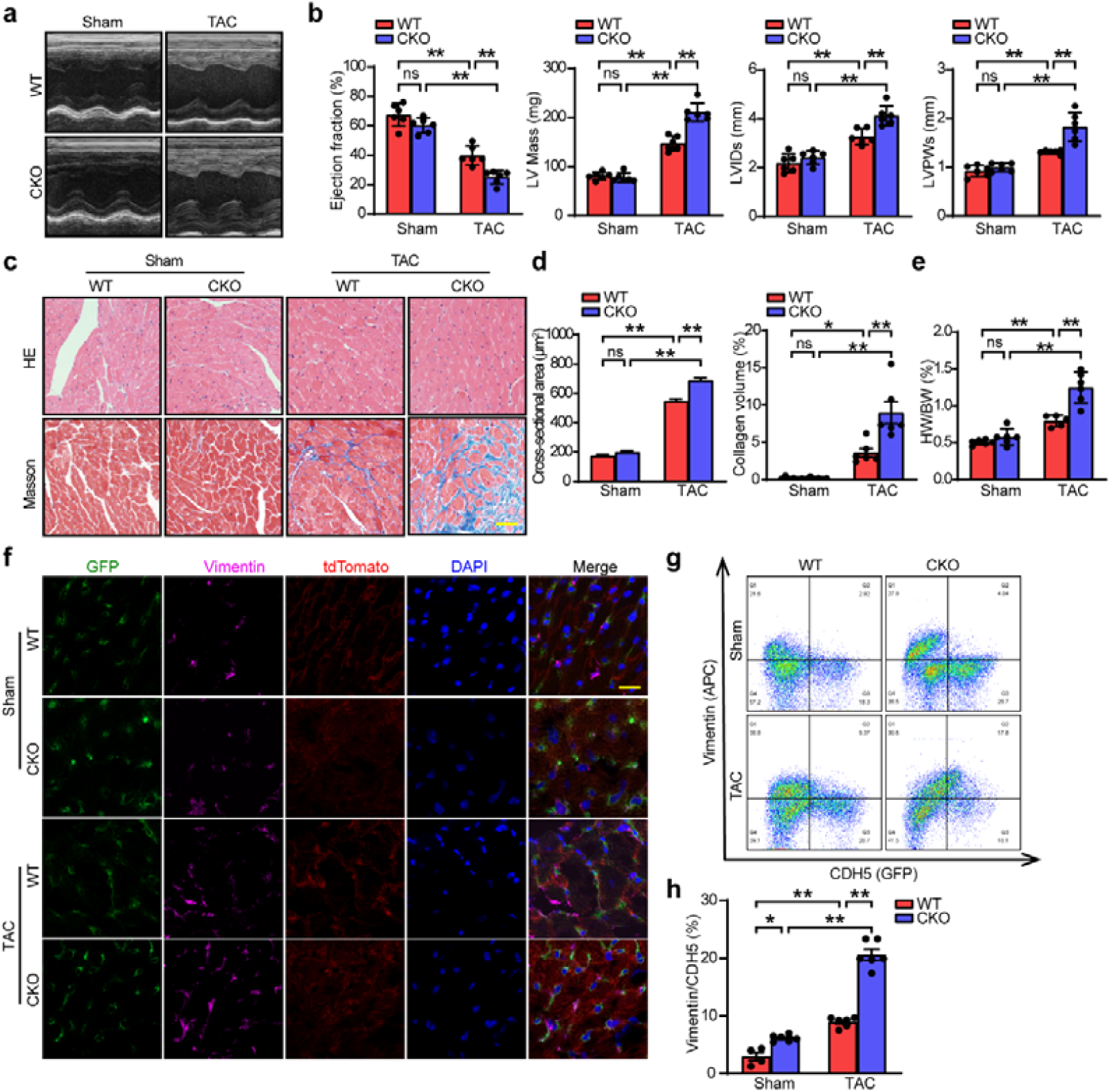
Endothelial cell-specific TET2 knockout aggravates cardiac function and fibrosis in pressure overloaded mice. (a-b) Echocardiographic measurements of CKO mice after TAC (n=6). (c) HE and Masson staining of CKO mice after TAC (n=5-6). (d) Statistical results of c. (e) Heart weight of body weight (n=5). (f) Immunofluorescence of WT and TET2-CKO hearts (n=6). (g) Flow cytometry of WT and TET2-CKO hearts (n=4-6). (h) Statistical results of g. Data are expressed as mean ± SEM. *P<0.05, **P<0.01.

Immunostaining of heart cryosections showed that vimentin co-localized with the GFP reporter, indicating that vimentin-positive cells were a subset of EndMT endothelial cells. In the TAC group, TET2-CKO mice exhibited more fibrosis and EndMT (Figure 3f), consistent with Masson staining. We further conducted a flow cytometry assay to identify how many ECs transdifferentiate into the mesenchymal cells. Vimentin positive cells in the TET2-CKO heart increased in both sham and TAC groups. Furthermore, the amount of EndMT cells in TET2-CKO mice was 2.3 folds higher than that of WT mice (p < 0.0001) (Figure 3g, 3h). These data suggested that endothelial TET2 regulated EndMT and pathological cardiac remodeling in vivo.

### 3.4 Transcriptional profiling and dynamic epigenetic regulation of HUVECs with TET2 knocked down

To determine the mechanism by which TET2 regulates EndMT, we performed RNA sequencing (RNA-seq) on siNC and siTET2 HUVECs treated with TGF-β1 and IL-1β (Figure 4a). First, the principal component analysis revealed clear differences between the two groups. The siTET2 group clustered from the siNC group, suggesting transcriptional heterogeneity (Figure 4b). Volcano Plot indicated genes being up-regulated and down-regulated in TET2 knocked down HUVECs compared to the control group (Figure 4c). The down-regulated differentially expressed genes (DEGs) highlighed genes with more significant and higher fold change in blue compared to up-regulated DEGs in red. We also checked the markers being expressed in endothelial cells (VCAM, CD34 and VEGFA) (Figure 4d) and mesenchymal cells (ACTA2, COL1A, and TGF-βI) using RNA-seq data (Figure 4e), to verify the quality of the sequencing data. The heatmap shows the top 20 up and down-regulated DEGs based on p-value (Figure 4f).

**Figure 4.**
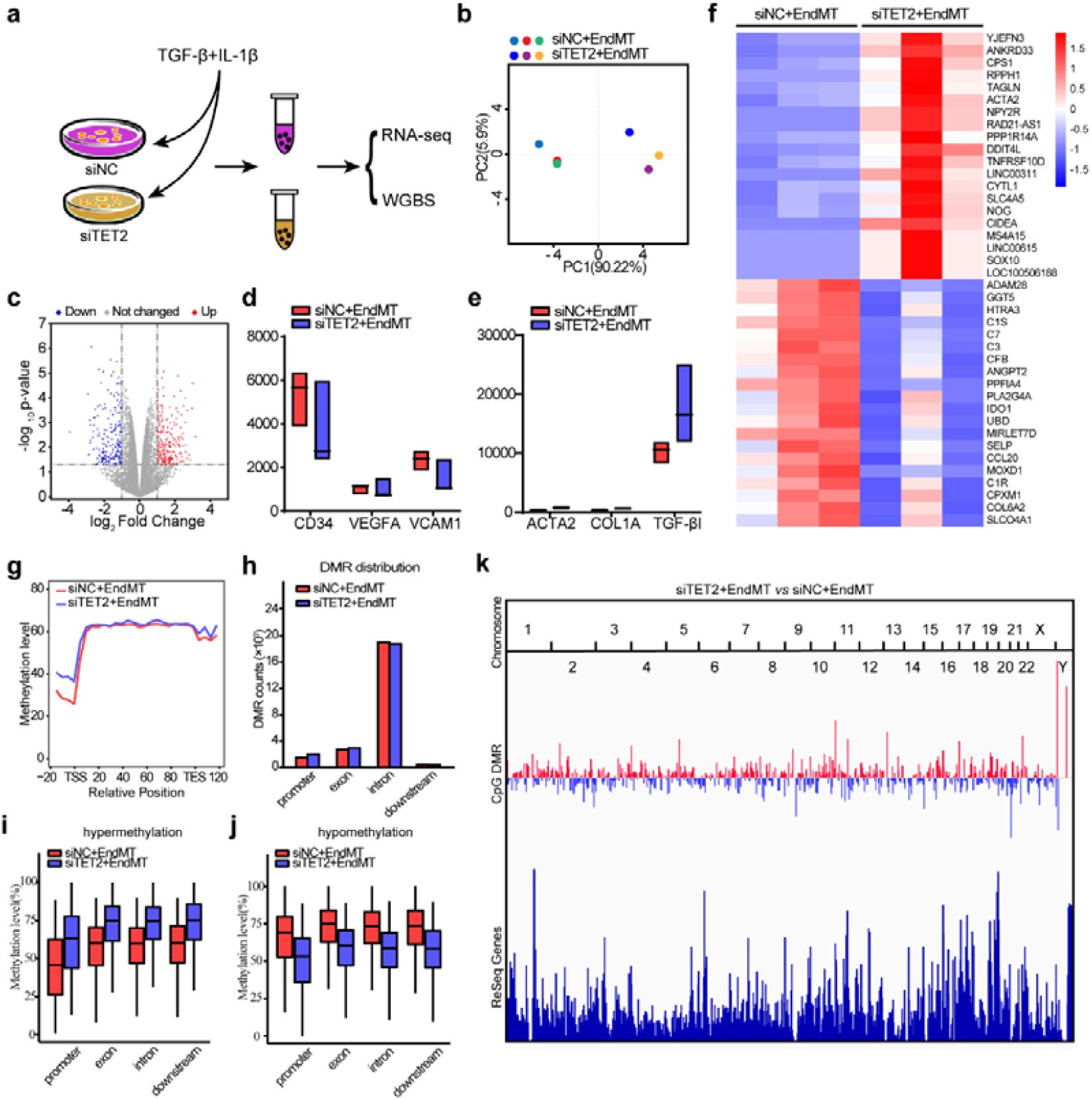
Transcriptional profiling and dynamic epigenetic regulation of HUVECs with TET2 knocked down. (a) Treatment of samples subjected to RNA sequencing and WGBS. (b) Principal component analysis of two groups. (c) Volcano Plot of two groups. (d-e) The expression of endothelial cells (VCAM, CD34, and VEGFA) and mesenchymal cells (ACTA2, COL1A, and TGF-βI) in the results of RNA-seq. (f) Heatmap of two groups. (g) Total DNA methylation of two groups. (h) The distribution of DMRs in different region. (i) DNA methylation levels of two groups in hypermethylation region. (j) DNA methylation levels of two groups in hypomethylation region. (k) The median methylation levels of DMRs.

Although the methylation levels were detected using ELISA and dot blot in our study before, we further sent the HUVECs being treated the same way in the above RNA-seq experiment for WGBS to understand the differences of DNA methylation after TET2 being knocked down in EndMT HUVECs. We observed that siTET2 HUVECs had higher CpG methylation levels (Figure 4g), particularly between the -20 hexamers and transcriptional start site (TSS), a promoter to which RNA polymerase bound. This provided clear evidence that TET2 could have worked as a demethylase to reduce DNA methylation. Based on the cutoff of DNA methylation percentage difference > 10% with p < 0.05, a total of 504,904 differentially methylated regions (DMRs) of CpG were obtained after comparing 8431 hyper-methylated and 5748 hypomethylated DMRs in the promoter region (Figure 4h). We also evaluated the DNA methylation in hypermethylation and hypomethylation regions (Figure 4i, 4j). SiTET2 HUVECs showed higher methylation levels in the hypermethylation region, while the control group showed higher methylation levels in the hypomethylation region.

We then analyzed significant changes in methylation on chromosome levels. The hypermethylated DMRs upon TET2 knockdown were plotted in red and hypomethylated DMRs were plotted in blue (Figure 4k). We found that hypermethylated DMRs concentrated on chromosomes 5, 10, 15, and Y, and the hypomethylated DMRs concentrated on chromosomes 2, 9, and 21. Since TET2 deficiency showed hypomethylated DMRs, therapeutics targeting downregulating TET2 could be an interesting direction for future study in these diseases.

We identified enrichment of the down-regulated DEGs (Suppl. Figure 9a) and up-regulated DEGs (Suppl. Figure 9b) into GO and KEGG pathways, and compared with pathways involving hypomethylated DMRs (Suppl. Figure 9c) and hypermethylated DMRs (Suppl. Figure 9d). The common pathways that DEGs and DMRs were mainly involved in plasma membrane biology, adding another layer of evidence supporting that TET2 participated in EndMT.

### 3.5 Loss of EGLN3 initiated EndMT in vitro as the downstream target of TET2

Considering TET2 as a demethylase, we then focused on down-regulated DEGs. DNA methylation of these genes was evaluated using WGBS data. The DNA methylation levels of down-regulated DEGs increased significnatly in siTET2 group, including promoter, exon, intron, and transcription termination site (TTS) (Figure 5a). After taking the intersection of down-regulated expressed genes in RNA-seq and up-regulated of promotor methylation levels in WGBS, we found 25 co-genes (Figure 5b). The top six genes were FRMD7, WDR17, TNFRSF9, MOXD1, EGLN3, and TTC23L according to the fold change of DEGs, DNA methylation difference of DMRs, and p-value (Suppl. Table 3).

**Figure 5.**
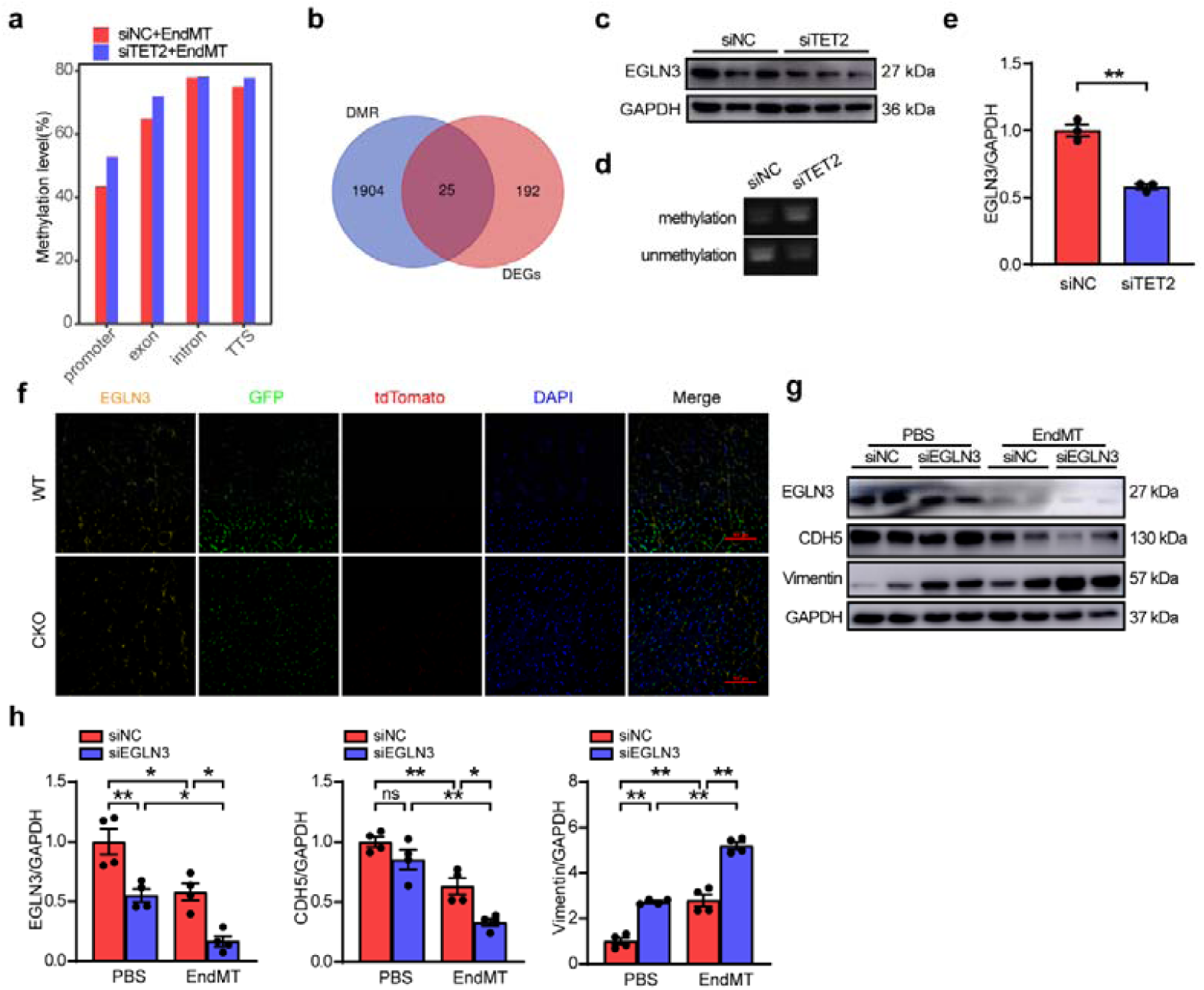
Loss of EGLN3 initiated EndMT in vitro as the downstream gene of TET2. (a) Methylation levels of down-regulated DEGs in different region. (b) Venn diagram of DEGs and DMRs. (c) The expression of EGLN3 after TET2 knocked down (n=3). (d) DNA methylation level of EGLN3 by MSP (n=4). (e) Statistical results of c. (f) Immunofluorescence of EGLN3 (n=6). (g) Western blot of HUVECs undergoing EndMT with or without EGLN3 knockdown (n=4). (h) Statistical results of g. Data are expressed as mean ± SEM. *P<0.05, **P<0.01.

We further explored the expression of these six genes in human tissue using The Human Protein Atlas (https://www.proteinatlas.org/) (Suppl. Figure 10), and found that EGLN3 was highly expressed in heart tissue. Thus, we investigated the role of EGLN3 in endothelial cells and explored its relationship with TET2. The protein level of EGLN3 was reduced in SiTET2 HUVECs (Figure 5c,5e). The methylation-specific PCR (MSP) also showed that the increased EGLN3 methylation could be blunted by knocking down TET2 (Figure 5d). In vivo, we used immunofluorescence to detect the levels of EGLN3 in the heart of TET2-CKO mice, confirming the finding from MSP (Figure 5f). We next knocked down EGLN3 in HUVECs and verified the role of EGLN3 in EndMT-modeling. In this HUVECs EndMT-model, the expression of EGLN3 decreased with the degree of EndMT being promoted upon EGLN3 knockdown (Figure 5j-5h), consistent with the result of RNA-seq. Together with the other’s finding that loss of EGLN3 mediated TGF-β/Smad signaling through TGFα44, these data indicated that EGLN3 might be the downstream gene of TET2 involved in EndMT.

### 3.6 Vitamin C alleviates myocardial remodeling by overexpressing TET2

Our above results indicated that deficiency of TET2 promoted EndMT, leading to myocardial fibrosis and cardiac dysfunction, we then tried to explore a practical approach to increase the expression of TET2 as a clinical application for preventing cardiac remodeling and fibrosis. Numerous reports have demonstrated that vitamin C could stimulate the catalytic activity of TET2^[22, 23]^. To further test the role of vitamin C on TET2 expression in endothelial cells, we added vitamin C to HUVECs and found that TET2 expression was remarkably elevated (Suppl. Figure 11). Therefore, we injected the WT and TET2-CKO mice with PBS or vitamin C intraperitoneally every 2 days after TAC for a total of 4 weeks (Figure 6a). We found the heart function of both WT and CKO mice injected with vitamin C were significantly improved compared to mice injected with PBS after TAC (Figure 6b, 6d-h). Notably, EndMT caused by the loss of TET2 almost blunted in WT mice after injection of vitamin C, and the number of endothelial cells transited to fibroblast in CKO also decreased after being treated with vitamin C (Figure 6i). The HE and Masson staining showed consistent results with the echocardiography (Figure 6c, 6j-6k). The decreased hypertrophic biomarkers, such as ANP and BNP, also supported the beneficial effect of vitamin C on myocardial remodeling (Figure 6l, 6m). The results demonstrated that vitamin C could alter the steady-state of DNA methylation during endothelial-mesenchymal transition in endothelial cells by enhancing TET2 activity and could be a promising candidate for the treatment of pathological cardiac remodeling in clinical practice (Figure 7).

**Figure 6.**
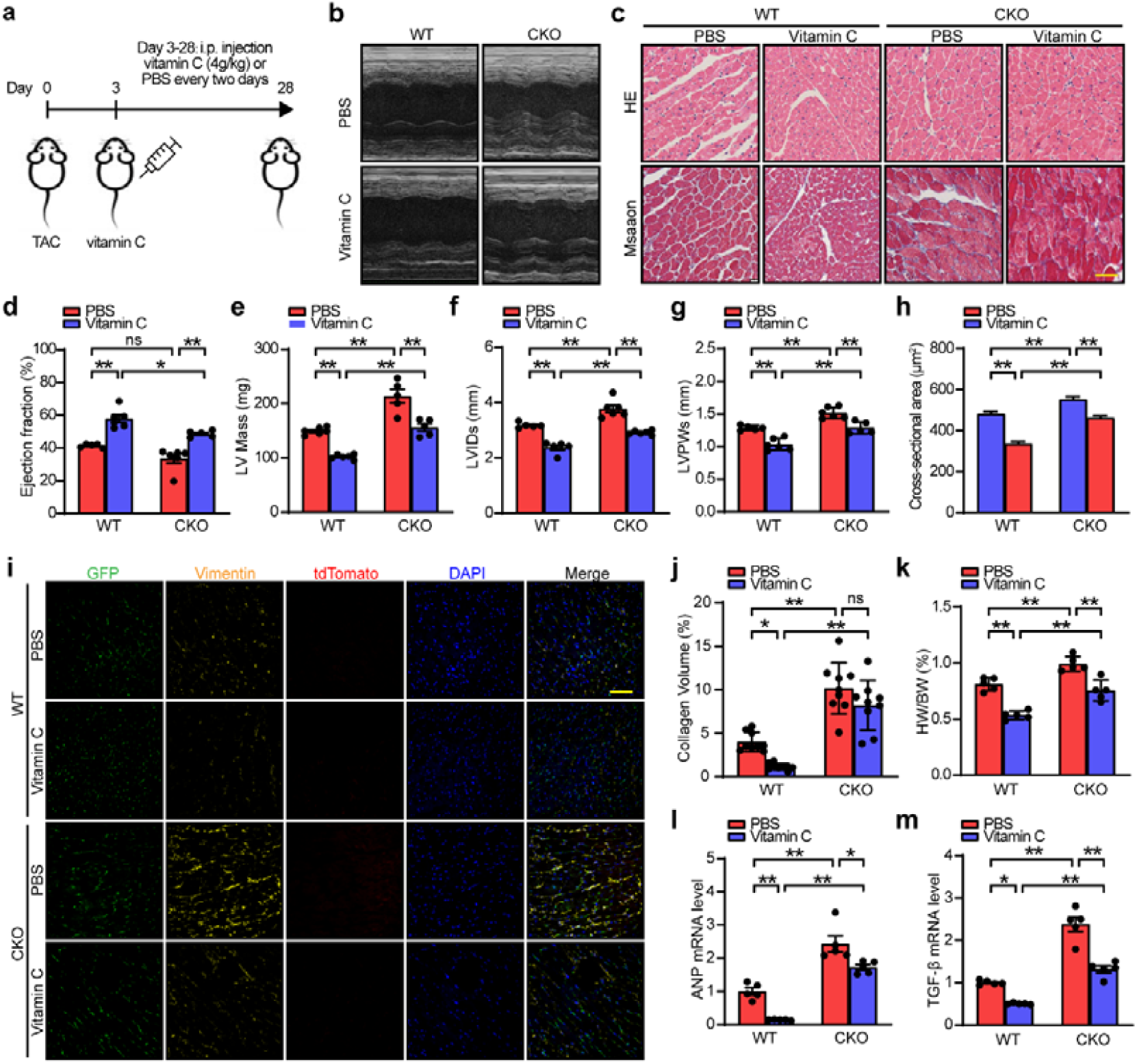
Vitamin C alleviates myocardial fibrosis induced by overexpressing TET2. (a) Treatment of mice. (b) Echocardiographic measurements. (c) HE and Masson staining. (d-g) Statistical results of b. (h) Statistical results of c. (i) Immunofluorescence of WT and TET2-CKO hearts after Vitamin C injection. (j) Statistical results of c. (k) Heart weight of body weight. (l-m) Relative ANP and BNP mRNA levels. Data are expressed as mean ± SEM. *P<0.05, **P<0.01.

**Figure 7.**
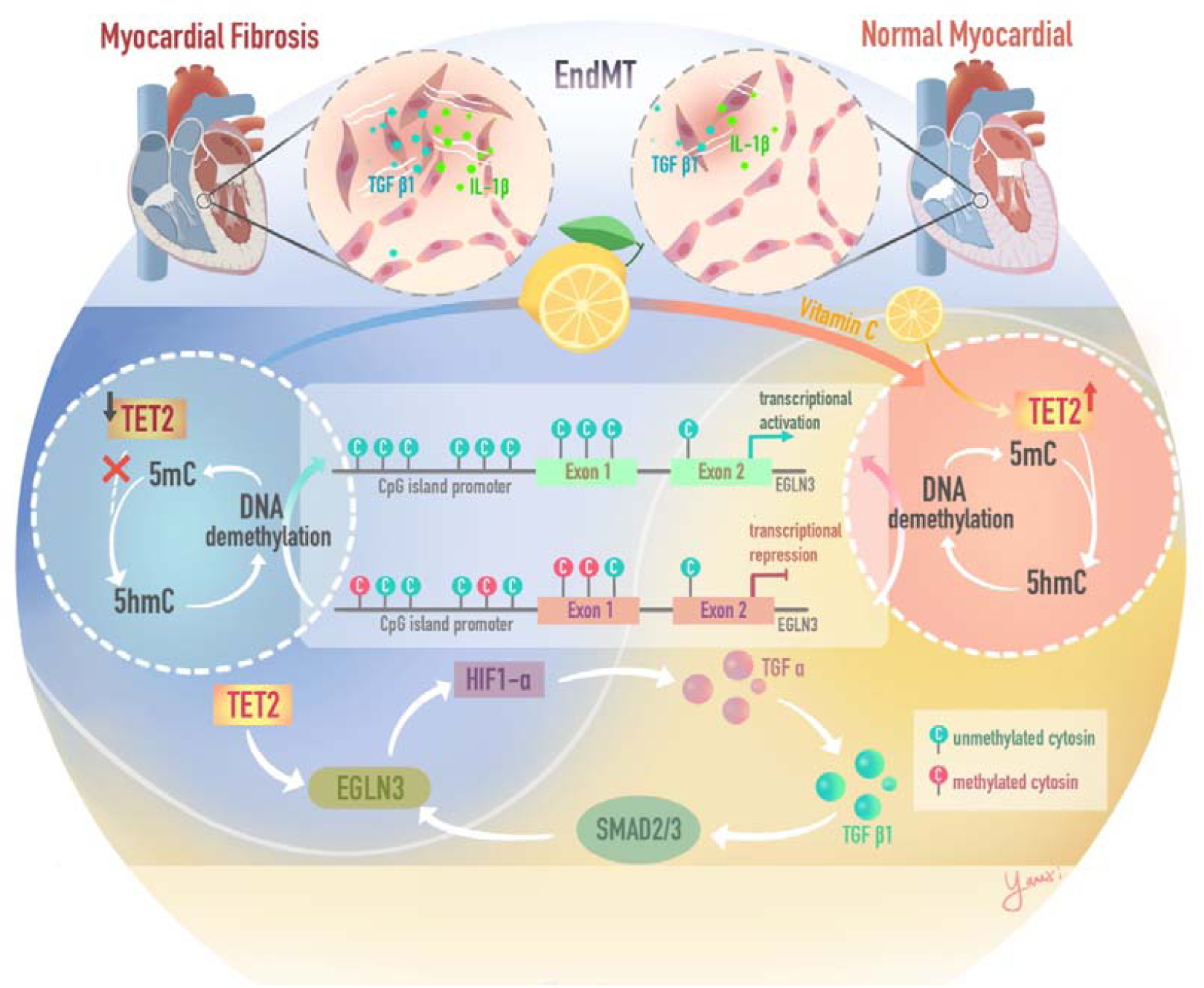
Mechanism diagram.

## 4. Discussion

Cardiovascular diseases are becoming a threat to human health worldwide^[24]^, particularly for an aging population nowadays. Heart disease is usually accompanied by cardiac remodeling and finally progresses to heart failure in which cardiac fibrosis participates as a synergistic process^[25]^. Accumulating evidence has suggested that epigenetic modification and regulation, such as DNA methylation, are involved in cardiovascular aging ^[26, 27]^. Our previous study demonstrated that DNA methylation participated in atherosclerosis^[12]^. In this study, we further explored the role of DNA methylation in cardiac remodeling. The application of DNA methylation inhibitor 5-aza alleviated myocardial hypertrophy and fibrosis in mice, indicating decreased methylation levels of DNA methylation could attenuate myocardial remodeling, and DNA demethylation might be a therapeutic approach for fibrosis.

Xiao et al. found that rats’ heart hypertrophy and failure would be rescued after being treated with the DNA methylation inhibitor 5-aza-2’-deoxycytidine^[28]^, consistent with our finding. Our data further suggested that DNA methylation mainly influenced cardiac fibrosis and myocardial interstitium. It should be noted that 5-aza primarily acts on the de novo DNA methyltransferases DNMT3a^[29]^. Madsen et al. reported that DNMT3a mediated cardiomyocyte metabolism and contractility by using human induced pluripotent stem cells^[30]^. Thus, we could not entirely exclude the role of DNMT3a on the protective role of 5-aza on cardiac remodeling in our experiments. The function of DNMT3a and DNMT3b is to maintain DNA methylation during cell division in cooperation with DNMT1^[31]^. However, DNA replication and cell division of cardiomyocytes were rarely happening in adult hearts. Thus, we rationalized that DNMT3a should play limited roles in our model. All the three TET family members (TET1, TET2, and TET3) can iteratively oxidize 5-methylcytosine (5-mC) into 5-hydroxymethylcytosine (5-hmC) and have similar structures, although TET2 seemed to have the most active on 5mC-DNA substrates. Xu x et al. reported that TET3 could demethylate the hypermethylation promoter region of RASAL1 and counteract the cardiac fibrosis effect stimulated by TGF-β^[32]^. Marco Sciacovelli finds that fumarate inhibits the demethylation of a regulatory region of mir-200ba429 mediated by TET2 and TET3, leading to enhanced migratory properties and the expression of EMT-related transcription factors in cancers^[33]^. Although multiple studies have revealed the role of different members of TET in a variety of cell types, endothelial cells mainly express TET2 rather than TET1 and TET3^[34]^.

The present study shows for the first time that the down-regulation of TET2 promoted cardiac remodeling by mediating endothelial to mesenchymal transition (EndMT). By using three different mouse models of myocardial remodeling, including TAC, diabetes, and MI, we found that the inhibition of fibrosis, not the cardiomyocyte, would influence cardiac remodeling and heart function. Inhibiting cardiac fibrosis is a two-edged sword in preventing heart failure. On the one hand, inhibiting cardiac fibrosis is necessary to maintain heart function. On the other hand, excessive inhibition of the fibrosis would lead to inefficient collagen production and ventricular rupture in myocardial infarction^[35]^. As such, post-injury cardiac fibrosis is a fine-tuned process requiring tightly-controlled mechanisms.

Although anti-fibrotic therapies have been proposed to be helpful in improving cardiac function in diseased hearts, the development of such therapies has been limited by an incomplete understanding of the origin of fibroblasts in the heart. Cardiac fibroblasts are composed of resident adventitial and interstitial fibroblasts^[36]^. Besides resident fibroblasts, injury-induced fibroblasts in hearts were reported to stem from a variety of cell origins, such as endothelial cells, bone marrow-derived fibrocytes or immune cells, mesenchymal stem/progenitor peri vascular cells, and adult epicardium ^[37]^. EndMT was first described by Leonard M Eisenberg in cardiac development in 1995 and proposed as a key mechanism for fibroblast generation after MI and experimental diabetes mellitus^[38]^. Yet, recent studies using Cdh5Cre or Tie2-Cre lineage tagging to label preexisting vascular endothelial cells failed to support that fibroblasts originated from endothelial cells in pressure overload model^[39]^.

Here, we showed that cardiac fibrosis was associated with the emergence of fibroblasts with an endothelial cell origin, and the deficiency of TET2 function plays a positive role in EndMT. The animal model we used in the current study was Cdh5-CreERT2^+/-^; Rosa26-mTmG^+/-^ mice. The transdifferentiation rate was about 10%, and the ECs in TET2-CKO mice would double the transdifferentiation rate. Although the transdifferentiation rate of ECs was not high, ECs acted on the fibroblast or ECs via paracrine. ECs secreted TGFβ that would stimulate the secretion of ECM from fibroblast. This could explain why the low transdifferentiation rate of EndMT had prominent cardiac fibrosis.

Using both WGBS and RNA-Seq, we investigated the potential mechanisms that may promote EndMT by maintaining DNA hypermethylation in the HUVECs with TET2 knockdown. So far, it is the only data set involving TET2 that include both WGBS and RNA-Seq on the same samples. The sequencing results indicated endothelial cell markers decreased and fibroblast markers increased on transcriptome level. Meanwhile, TET2 knockdown HUVECs had more methylated promotors on CpG, which was consistent with our previous experimental results. Besides, common pathways are mainly concentrated on cellular components. When EndMT occured, endothelial cells lost cellular delaminate and adhesion, and reorganized their cytoskeleton to form spindle-shape cells^[40]^. Our research showed that collagen and cytokines like TGF-β release increased in siTET2 HUVECs. The change could also affect the cardiac microenvironment by activating paracrine signals^[41]^, which worsen heart dysfunction. This may explain why the cross-section area of myocytes was larger in TET2-CKO mice.

In order to determine whether DMRs correlated with changes in gene expression, we take the intersection of up-regulated DMR-associated genes and down-regulated DEGs. This comparison revealed a total of 25 co-genes. Based on the fold change of DEGs, DNA methylation percentage difference of DMRs, p-value, and expression levels in heart tissues, EGLN3 was found to be the downstream gene candidate. EGLN3 is a member of alpha-ketoglutarate-dependent dioxygenases that regulate hypoxia-inducible factor (HIF) by prolyl hydroxylation^[42, 43]^. We confirmed that the expression of EGLN3 decreased in TET2 defected HUVECs in vitro and the heart of TET2-CKO mice in vivo, proving that TET2 worked as an upstream gene to regulate EGLN3. In addition, we demonstrated that the methylation level of EGLN3 increased after TET2 knockout by MSP. Furthermore, the expression of EGLN3 could suppress EndMT in cardiovascular fibrosis ^[44, 45]^

Although Renin-Angiotensin System (RAS) inhibitors and β blockers had proved effective in preventing cardiac fibrosis and remodeling by antagonizing the RAS systems, specific therapies to restrain or reverse cardiac fibrosis have not been available yet^[46]^. Thus, exploring the underlying mechanisms of cardiac fibrosis and developing therapeutic targets is urgently required. Our study provides a new molecular mechanism for the role of TET2 in EndMT-induced cardiac fibrosis. It provides further evidence for targeting TET2 as a potential therapeutic option for alleviating cardiac fibrosis and preserving cardiac function in patients with cardiac hypertrophic and remodeling.

Multiple lines of evidence have suggested that vitamin C could promote TET2 activity and enhance TET2-mediated DNA demethylation^[47]^. Our results demonstrated that vitamin C enhanced the activity of TET2 in HUVECs. Vitamin C could improve cardiac function and alleviate myocardial remodeling when systematically injected into mice. Restoration of TET2 activity after transverse aortic constriction blocked the occurrence of EndMT in mice by high-dose vitamin C treatment. Put all together, the beneficial role of vitamin C in preventing heart function has stemmed from multiple lines of reasoning and pieces of evidence. Interrupting EndMT caused by the loss of TET2-mediated demethylation could also be a unique therapeutic approach to treating various fibrotic diseases.

A short course of intravenous vitamin C in pharmacological dose seemed to be a promising, well-tolerated, and cheap adjuvant therapy to modulate the overwhelming oxidative stress in the initial stage of myocardial fibrosis. Large randomized controlled trials are necessary to provide more evidence in the future.

## Conclusions

In this study, we found that dysfunction of TET2 in endothelial cells accelerated cardiac remodeling by regulating EndMT. The activation of TET2 counteracted cardiac remodeling and improved heart function. Besides, our research screened EGLN3 as the potential targets of TET2 by DNA methylation sequencing and studied the related signaling pathway. Finally, vitamin C which enhanced TET2-dependent DNA demethylation could prevent left ventricular remodeling after TAC in mice and improve cardiac function.

## Acknowledgments

We thank patients who participated in this study. This study is supported by grants No. 82070230, 91939101 and 81700210 from the Chinese National Natural Science Foundation, Clinical Research Plan of SHDC (No. SHDC2020CR4019), and Clinical Research Plan of Shanghai’s health commission (No.20214Y0152)

## Author Contributions

Wenxin Kou performed the cell function, animal experiments, and conducted western blot, immunofluorescence, and analyzed data. Yefei Shi performed the cell culture, animal experiments and western blot. Bo Li performed the animal experiments and flow cytometry. Yanxi Zeng drew mechanism diagram. Qing Yu and Xinyu Weng performed TAC model. Shiyu Gong performed MI model. Ming Zhai and Shuagjie You performed the bioinformation analysis. Jianhui Zhuang and Yifan Zhao conducted DNA methylation experiments and interpreted results. Weixia Jian and Yawei Xu conceived the project. Wenhui Peng conceived the project, designed experiments, analyzed data, interpreted results and wrote the manuscript.

